# Tapestry: validate and edit small eukaryotic genome assemblies with long reads

**DOI:** 10.1101/2020.04.24.059402

**Authors:** John W. Davey, Seth J. Davis, Jeremy C. Mottram, Peter D. Ashton

## Abstract

**Summary:** Small eukaryotic genome assemblies based on long reads are often close to complete, but still require validation and editing. Tapestry produces an interactive report which can be used to validate, sort and filter the contigs in a raw genome assembly, taking into account GC content, telomeres, read depths, contig alignments and read alignments. The report can be shared with collaborators and included as supplemental material in publications.

**Availability:** Source code is freely available at https://github.com/johnomics/tapestry. Package is freely available in Bioconda (https://anaconda.org/bioconda/tapestry).

---

Long read sequencing technologies such as Oxford Nanopore and Pacific Biosciences can now produce reads spanning many repetitive sequences in eukaryotic genomes. This means raw genome assemblies based on long reads can be close to complete, particularly for small genomes, in contrast to assemblies of read sets from older sequencers which would typically contain thousands to hundreds of thousands of contigs (contiguous sequences). However, further processing is usually required to turn a raw long read assembly into a complete, validated assembly. Raw contigs must be classified as complete chromosomes, accessory genomes, or fragments that have not assembled completely, and it may be possible to explain why fragments have failed to assemble into complete chromosomes. Many different aspects of the assembly, such as presence of telomeres, GC content, circularity, and alignments of reads and other contigs, can be used to validate contigs. Tapestry is a tool to draw these features together to allow the manual labelling and filtering of a raw eukaryotic genome assembly. It is intended for close-to-complete assemblies of small eukaryotic genomes (<50 megabases (Mb)) with less than a few hundred contigs, but can run on larger genomes with some limitations (see Discussion).

Tapestry is intended to complement other tools such as QUAST (Mikheenko *et al.*, 2018), which will produce global assembly metrics and allow comparison to other assemblies, IGV (Thorvaldsdóttir *et al.*, 2013), which will allow close inspection of read alignments, and BUSCO (Waterhouse *et al.*, 2018) or FGMP (Cissé & Stajich, 2019), which assess gene set completeness. It is a genome assembly assessment tool similar to Knot (Marijon *et al.*, 2019), Bandage (Wick *et al.*, 2015) and Merqury (Rhie *et al.*, 2020), but allows inspection of eukaryotic features and read alignments, allows assessment and filtering of individual contigs, and can be used for assemblies where no graph is available.

Tapestry generates a HTML report that can be loaded into modern web browsers. The report presents a visualization of all contigs in the assembly, reporting contig size, GC content, and read depth in a table, and visualizing telomeres, ploidy estimates, contig alignments and read alignments. The report is interactive; contigs can be sorted, filtered and labelled within the HTML report, and a final set of contigs can be exported as a CSV file. This CSV file can be used to produce a filtered assembly and can also be loaded back into the report to demonstrate the filtering process. The HTML report and CSV file are self-contained files that can be shared with collaborators or included in publications, without requiring a web server or installed software beyond a JavaScript-enabled web browser.

Tapestry contains two tools, *weave*, which will generate a report on a raw assembly, and *clean*, which will filter the assembly based on information exported from the report. *weave* accepts a raw assembly in FASTA format, a set of reads in FASTQ format (which may be gzipped), and an optional telomere sequence, and produces a HTML report about the assembly and some additional analysis files, including a TSV file of contig statistics and alignment BAM files. It subsamples the reads to a user-specified depth (default 50x), using a user-specified minimum read length (default 10,000); reads are filtered because not all reads are needed to estimate Tapestry’s read depth features, and short reads that do not map uniquely are unlikely to be useful for validation, as they may align inaccurately to repetitive sequences. Reads and assembly contigs are then aligned to the assembly using minimap2 (Li, 2018). Tapestry then calculates a range of summary statistics for each contig and builds a report containing the summary statistics and the read and contig alignments for all contigs. *clean* takes the raw assembly in FASTA format and a list of contigs exported from the Tapestry report in CSV format and produce a cleaned assembly FASTA. Further details on implementation, calculation of summary statistics, and use of these statistics for assembly validation, can be found in Supplementary Information.

To demonstrate Tapestry, we generated a version of the complete *Cyanidioschyzon merolae* genome (20 chromosomes, 16.5 Mb long, plus mitochondrial (32 kb) and chloroplast (150 kb) genomes; Nozaki *et al*., 2007) containing variants simulated with simuG (Yue and Liti, 2019), simulated a long read set of the genome with badread (Wick, 2019), assembled the reads, and produced a Tapestry report for the assembly (see Supplemental Information, Genome Simulation). The variants included 1 translocation between chromosomes 17 and 19, which was intended to produce incomplete chromosome fragments, 20 100bp-100kb copy number variants, which may generate haplotype contigs (for long deletions or tandem duplications) or broken chromosomes (for dispersed duplications), and an extra haplotype of chromosome 3 to simulate a triploid chromosome. A read set of 236,876 reads, 3.4 GB in size, was simulated for this genome and assembled with canu (Koren *et al.*, 2017) to produce a 66 contig assembly, 18.9 Mb long. Tapestry produced a report for this assembly in 10 minutes on a 2012 MacBook Pro using 4 cores and maximum 4 GB RAM.

The *C. merolae* Tapestry report (Supplementary File 1) was manually annotated with reference to the published *C. merolae* genome (Supplementary File 2, which can be loaded into Supplementary File 1). The report shows that the assembly contains 16 complete nuclear chromosomes, 4 fragmented or incomplete nuclear chromosomes in 7 contigs, complete mitochondrial and chloroplast genomes, and 41 additional contigs. While the assembly has been annotated using the reference, these features are identifiable without the reference sequence (see Supplementary Information). The complete chromosomes are shown by the presence of two telomeres, consistent nuclear diploid read depth of ∼40, GC content of ∼54% and no reads aligning beyond the ends of contigs (see, for example, tig00000083 in the report). tig00000092 has increased read depth of 58, suggesting triploidy; this is the assembly of chromosome 3 which has an extra simulated haplotype. The mitochondrial (tig00000004) and chloroplast (tig00000091) contigs are easily identifiable by GC content and read depth, and by their lack of alignments to any other contig except themselves (visible by clicking the names of these contigs in the plot to show contig alignments); their self-alignments at either end suggest they are circular and the contigs need to be trimmed of redundant material.

Two incomplete contigs, tig00000094 and tig00003757, overlap with each other at their incomplete ends, and tig00003757 has a set of reads with overhangs that align to tig00000094 (visible by hovering over purple overhangs in the read alignment plot for tig00003757, selected by the radio button for tig000003757 in the report’s contig table). These two contigs are two halves of chromosome 14; a deletion in one haplotype at the broken region may have prevented complete assembly of the chromosome. One other incomplete contig, tig00000020, has an alignment to the middle of complete chromosome tig00003747 and also to incomplete contig tig00000089. These are caused by the translocation between contigs 17 and 19. Although confirming and validating the breakage of chromosome 14 and the translocation would require more analysis beyond that provided in the Tapestry report, the report does allow the contigs to be linked by alignments and labelled for further investigation.

Of the 41 additional contigs, 23 can be identified as haplotypes caused by copy number variants, or extra copies of subtelomeric sequence. These can be identified by haploid or lower read coverage, and contig alignments to complete chromosomes. The remaining 18 contigs have very low read depth, no alignments to any other contig, are relatively short, and many have extreme GC contents. They can be labelled as junk or unknown contigs and removed from the assembly. These contigs were likely assembled from simulated reads with low-complexity or random sequence.

The final genome annotation (Supplementary File 2) was exported from the Tapestry report and could be passed to *clean* to produce a filtered FASTA file with 24 contigs: 16 complete chromosomes, 4 incomplete chromosomes in 6 contigs and two accessory genomes.

Tapestry is designed for manual clean up of small genomes (<50 Mb) with few contigs (<500), but it will run on larger genomes. To do this, it does not output read alignments to the report, as they make the report too large to view in a normal web browser (though read alignments can be forced if necessary). However, it still uses read alignments to calculate read depth statistics. The report and TSV files can still be used to explore the genome assembly. Tapestry generated a report for the human GRCh38 primary assembly (3.1 Gb, 194 contigs) with simulated reads (10x coverage of the genome) in 4 hours on a 32-core server with 32 GB of RAM.

While genome assembly algorithms are improving all the time, their output is hard to understand for users new to assemblies and often requires bioinformatics skills. We hope that Tapestry will make genome assemblies easier to understand and complete for all users.

## Supporting information

Supplementary File 1: Tapestry report

Supplementary File 2: Genome annotation

