## Supplementary Information for "Tapestry: validate and edit small eukaryotic genome assemblies with long reads"

### Supplementary Methods

Tapestry is written in Python. Reads are aligned to the raw assembly using minimap2 in mode map-ont. The assembly contigs are then self-aligned using minimap2 in mode asm20 with options -D, -P and setting -m and -s to a user-specified minimum contig alignment length (default 2,000 bp). Tapestry stores alignments in an SQLite3 database (https://sqlite.org/index.html) with SQLalchemy (https://www.sqlalchemy.org) and builds its report using Jinja (https://jinja.palletsprojects.com/en/2.11.x/) and D3 (https://d3js.org).

Tapestry generates the following information for each contig (details explained below):
- length in basepairs
- GC content
- presence of telomeres
- read alignments
- read depth
- ploidy estimates
- contig alignments

Telomere sequences are counted in the first and last 1000 bp of each contig, based on a telomere sequence provided by the user. The Tapestry report shows telomeres as red vertical lines at the ends of contigs, with lighter lines representing fewer copies of the telomere sequence.

Tapestry processes minimap2 read alignments for each contig by calculating the lengths of the original read clipped from the left and right of the aligned region of the read, searching for reads with multiple alignments, and identifying contigs that are connected by reads with multiple alignments. This information is reported in a read alignment plot for each contig in the Tapestry report. By default, read alignments are only output for assemblies smaller than 50 Mb, but users can force output of read alignments for larger assemblies with option -f.

Read depth is calculated for each window across the genome. Window size can be supplied by the user but by default is set to ~1/30th of the assembly's contig N50 length. The window read depth is calculated by summing all bases aligning within the window and dividing by window size. The contig read depth reported by Tapestry is the median window read depth across all windows for the contig.

Ploidy estimates for each window across each contig are generated by calculating the median window read depth across the entire assembly and assuming that this read depth represents diploid read depth. Tapestry then calculates estimated haploid, diploid, triploid, tetraploid and repeat read depths based on the median read depth, and assigns each window to a ploidy by calculating which ploidy depth estimate the window depth is closest to. The window ploidy estimates are included in the Tapestry report by colouring the contig diagrams by ploidy level, and also showing ploidy levels in the read alignments plot.

Contig alignments from minimap2 are also included in the Tapestry report and can be shown by clicking on the contig name in the diagram.

These features can be used to classify and filter contigs, as described in the following section.

### Feature Identification

Tapestry is designed to assist the identification of complete chromosomes from the nuclear genome of interest, accessory genomes, chromosome fragments, haplotypes, complex structural variations, and extraneous contigs such as extra repeat or subtelomere copies. This section explains how contigs can be classified into these categories using Tapestry's report.

**Complete chromosomes**
If a raw contig is a complete chromosome, it should have following features:

- **Telomeres** should appear at both ends. Tapestry identifies these by asking the user for a telomere sequence to search for and then counting the number of telomere sequences in the first and last 1kb of the contig. In the report, telomere presence is shown by vertical lines at the ends of contigs, with lighter lines representing fewer telomere copies found (with maximum opacity for 20 telomere copies).

- **GC content** of the contig should be relatively similar to GC content of other nuclear contigs. Tapestry reports GC content of each contig in a table and allows contigs to be sorted by GC content so they can be clustered.
- The depth of read alignments to the contig should be relatively consistent and close to the median read depth for all nuclear contigs. For example, if the genome has been sequenced 20 times, and the contig is complete with no major errors, when the read set is aligned back to the genome there should be roughly 20 reads aligning to any part of the contig. Tapestry reports **read depths** in a table and reports **ploidy estimates** (explained above) visually so contigs can be classified accordingly.
- Reads should end at the end of the contig with no long overhangs. If the contig is a complete chromosome, there is no more DNA to sequence beyond the telomere, and therefore reads should not extend beyond the end of the contig. Conversely, if the contig is incomplete it is likely reads will extend beyond the end of the contig, as they represent some complex sequence the assembler was unable to complete. Tapestry visualizes read alignments so read overhangs beyond the end of a contig can be examined.

**Accessory genomes and contamination**

Accessory genomes such as mitochondria, chloroplasts or symbionts, or contigs from contaminating genomes, can usually be identified by variations in GC content and read depth from the nuclear genome (Challis *et al.*). These will be shown in the Tapestry report. Accessory genomes are also often circular; this sometimes results in accessory contigs containing identical overlapping sequence at their starts and ends, which can be visualized in Tapestry by examining **contig self-alignments**.

**Chromosome fragments**

If a contig is not a complete chromosome, but contains some unique region of the genome that could not be assembled completely, then reads are likely to align poorly to the ends of the contig and may have other alignments to other contigs, which may help to explain why the assembly is incomplete. The contig may also have overlapping regions aligning to other contigs. Tapestry can visualize these read and contig alignments so they can be manually inspected.

**Haplotypes**

Many genome assemblers aim to produce a representative haploid assembly for organisms with multiploid genomes. This means they must strike a balance between collapsing minor variations such as SNPs or indels and separating similar material from different chromosomes. This balance typically means that some regions of the genome with substantial haplotypic variation are reported as separate contigs, and not collapsed into one contig. These contigs can be identified in Tapestry because they usually align completely to some other contig, and the read depth for the aligned region is around half of the expected diploid read depth (for diploid organisms).

**Structural variation**

Features of multiploid genomes such as translocations, duplications, inversions and ploidy variations are usually not handled by genome assemblers and need to be identified after assembly. Tapestry can help to identify contigs involved in such variations by examining read and contig alignments and variations in read depth. Groups of contigs that are connected by alignments and have depth variations can then be reported together and investigated outside of Tapestry.

**Extraneous contigs**

Small contigs that align to many other contigs and have many or few reads aligning to them are usually extra copies of repeats or subtelomere sequences; some contigs may also be generated by bad quality raw reads, such as reads from blocked pores in Oxford Nanopore read sets. Tapestry can be used to identify these contigs and filter them from the genome.

### Genome Simulation

The *Cyanidioschyzon merolae* genome was downloaded from Ensembl Plants (accession ASM9120v1, toplevel DNA sequence). simuG v1.0.0 was used to add one translocation, 20 copy number variations (deletions and tandem and dispersed duplications between 100 and 100,000 bp long, roughly one for each chromosome), 16500 SNPs (1 in 1000 bases) and 1650 indels (1 in 10,000 bases) to the nuclear genome. Telomeres CCCCCCATT/AATGGGGGG are known to be present at all chromosome ends in tandem arrays 400-700bp long (Nozaki *et al.* 2007) but the assembly only includes 0-2 copies of the telomere sequence, so arrays of a random length between 400 and 700bp were simulated and concatenated to chromosome ends for detection by Tapestry. An extra copy of chromosome 3 (482kb long) was simulated with 1 tandem duplication, 482 SNPs and 48 indels, to simulate a triploid chromosome. The reference genome and simulated genome sequences were then merged together to make a complete example genome of 43 sequences (41 nuclear haplotypes, plus mitochondrial and chloroplast genomes).

badread v0.1.5 was used to simulate 50x coverage of the example genome (100x coverage of the diploid genome) in Oxford Nanopore reads (default parameters except no adapter sequences were ligated; all contigs assigned depth 1 except mitochondrion (depth 200) and chloroplast (depth 30), which were both labelled as circular). The simulated read set was then assembled with canu 1.9 using a genomeSize parameter of 16.7m and -fast mode. A Tapestry 1.0.0 report was generated for the canu assembly using the badread read set and a telomere sequence AATGGGGGG.
