## Supplementary File 1: Tapestry report for "Tapestry: validate and edit small eukaryotic genome assemblies with long reads": Supplementary_File_1_c_merolae_simulated.tapestry_report.html


##### JavaScript Disabled

Tapestry uses JavaScript, but it looks like
you have JavaScript disabled in your web browser.
Please enable JavaScript to view the full report.


Tapestry Report


- Assembly
- Contigs
- Reads
- Information

  

This is a Tapestry report on the genome assembly **c\_merolae\_simulated.badread\_50x.canu1.9.fasta**. See Information for full details.

If you have a contig order in a Tapestry CSV file, load it here:

Contigs in CSV file do not match contigs in report. Please choose a CSV file that matches this assembly.
×

Contigs loaded successfully!
×

---

### Assembly

Selected and rejected contigs are those checked and unchecked in the contig table **Keep** column below.

| Contig set | Contig number | Length (bp) | N50 number | N50 length |
| --- | --- | --- | --- | --- |

---

### Contigs

Red vertical lines at the ends of contigs show telomeres. Opacity of red lines reflects the number of telomere sequences found at each contig end. Green blocks show ploidy levels; see read alignment plot for key.

Click on a table radio button to show read alignments for the selected contig.

Click on plot contig names to show contig alignments. Hover over blue alignment blocks to show connecting alignments. The selected contig is shown with a dashed black line. Purple lines show contigs with alignments to the selected contig. The opacity of each purple line shows the percentage of the selected contig found in the aligned contig. Dark lines suggest the selected contig is mostly contained in the aligned contig.

|  | Contig | Length | GC % | Read Depth | Group | Keep | Notes |
| --- | --- | --- | --- | --- | --- | --- | --- |

---

### Read Alignments

The contig selected by radio button in the contig table is shown.

Read alignments are shown in blue.

Clipped parts of reads with no alignments are shown in beige.

Where parts of reads align elsewhere, lengths of unaligned read sequence connecting to neighbouring alignments are shown for alignments to the   
same contig or different contigs.

Hover over alignments to show names of connecting contigs (or '-' if none).

Alignment opacity shows mapping quality; darker alignments are higher quality.

---

### Information

| Option | Value/th> |
| --- | --- |

© 2019-2020 John Davey.
Built with D3,
Bootstrap,
Bootstrap Table,
X-editable,
TableExport,
FileSaver.js,
JQuery,
PapaParse,
Popper.js, and the
Litera theme via
Epic Bootstrap.
